# ExoFILT: Transfer learning for robust and accelerated analysis of exocytosis single-particle tracking data

**DOI:** 10.64898/2026.02.27.708581

**Authors:** Eric Kramer, Laura I. Betancur, Sasha Meek, Sébastien Tosi, Carlo Manzo, Baldo Oliva, Oriol Gallego

**Author notes:** Corresponding author. Department of Medicine and Life Sciences (MELIS), Universitat Pompeu Fabra (UPF), Barcelona 08003, Spain. (O.G.).

## Abstract

**Motivation:** Understanding constitutive exocytosis at the molecular level requires quantitative characterization of protein dynamics during the process. Single-particle tracking allows the measurement of protein dynamics in living cells. However, identifying bona fide exocytic events requires extensive manual annotation, limiting throughput and introducing personal biases that affect reproducibility.

**Results:** We present ExoFILT, a deep learning-based classifier designed to identify exocytic events in single-particle tracking data, using the exocyst complex as a reference. Trained via transfer learning on simulated and experimental data, ExoFILT reduces the time required for manual annotation by ten-fold while improving measurement consistency across researchers. When applied to simultaneous dual-color time-lapse movies, ExoFILT enabled the systematic quantification of temporal relationships between exocytic proteins. The increased throughput uncovered distinct subpopulations of exocytic events with differential molecular composition (e.g., events with and without detectable levels of Sec1), underscoring the potential of ExoFILT to reveal mechanistic insights into exocytosis.

## 1. Introduction

Constitutive exocytosis is a vesicle trafficking pathway that delivers cargo to the plasma membrane and the extracellular space. Unlike regulated exocytosis, which is characteristic of a few specialized cell types only, constitutive exocytosis (hereafter exocytosis) is essential for all eukaryotic cells. This secretory pathway was originally described in *Saccharomyces cerevisiae*, where more than 30 key components of exocytosis have been identified^1–7^. The genetic tractability of *S. cerevisiae* has consolidated it as the main model to study the molecular mechanisms of exocytosis, while the high conservation of the exocytic machinery across eukaryotes has allowed findings in yeast to be extended to other organisms.

The exocyst, a heterooctameric complex, plays a central role in exocytosis where it participates in both vesicle tethering and fusion steps^8,9^. The exocyst function involves the interaction with additional proteins, such as Sec1, a Sec1/Munc18 (SM) protein that plays a critical role in the assembly of the SNAREs (soluble *N*-ethylmaleimide-sensitive factor attachment protein receptors)^10^. Major advances have been made in elucidating the structure of the main components of the exocytic machinery, such as the exocyst^6,11–15^ and SM proteins^16–18^. However, a detailed mechanistic understanding of this process remains elusive because we still lack knowledge of the temporal organization of the proteins involved. Such temporal coordination is essential for resolving the sequence of molecular events that ultimately result in successful exocytosis.

When expressed as fusions with fluorescent proteins, the exocyst and other components of the exocytic machinery can be imaged as diffraction-limited fluorescent puncta at the plasma membrane^19^. Single-particle tracking (SPT)^20,21^ is a live-cell imaging technique that allows the detection and linking of such puncta along the frames of time-lapse movies, capturing their temporal behaviour along the process of exocytosis^8,19,22–24^. Quantitative analysis of the obtained tracks can measure biologically relevant features, such as kinetic parameters (e.g., diffusion coefficients), molecular copy number, or spatiotemporal dynamics^8,19,23–27^, providing insights into the functions and mechanisms of the proteins under study. Unfortunately, SPT of exocytosis is limited by biological and technical factors that challenge the systematic detection of exocytic events and prevent comparative analysis between different researchers.

The polarized nature of exocytosis combined with the size of secretory vesicles (∼80 nm in diameter^8^, smaller than the diffraction limit), hampers the ability to spatially resolve neighboring exocytic events. The copy number of proteins participating in each exocytic event is small (e.g., 7 exocysts control each exocytic event on average, in a 1:1 stoichiometry with other key proteins)^8^, which reduces the signal-to-noise ratio (SNR). Thus, state-of-the-art particle tracking algorithms often fail in accurately capturing bona fide exocytic events, even with the aid of parametric filters^8,24^. Therefore, SPT results in substantial false positive tracks that need additional manual inspection, a step that requires highly specialized researchers, is time-consuming and is prone to personal biases. These technical limitations are not unique to the analysis of exocytosis but reflect a broader challenge for the SPT community. Signal is often limiting because light toxicity and photobleaching require low illumination to maximize the length and framerate of the time-lapse movies. This highlights the need for computational frameworks that can improve robustness and efficiency in low SNR regimes.

Methods based on mathematical modeling^28–32^, and more recently, deep learning (DL)^33–35^, have shown promising performance in imaging constitutive exocytosis in mammalian cells. Nearly all of these tools leverage the properties of the pH-sensitive fluorophore pHluorin or related variants, which produce a distinctive high SNR fluorescence burst upon vesicle fusion. Unfortunately, pHluorin-based methods do not perform well in yeast cells because of poor folding, secretion and maturation of the probe^36–38^. Moreover, pHluorin reflects only local pH changes associated with fusion pore opening, which cannot be leveraged to follow the dynamics of labelled proteins during the rest of the pathway. Similarly, pHluorin does not capture the complete panel of physiological contexts associated with exocytic events, as it fails to label those exocytic events that do not undergo vesicle fusion, which hides functional and mechanistic insight. Consequently, the direct observation and quantification of the dynamics of the exocytic machinery remains challenging.

To overcome these limitations, we developed ExoFILT (Exocytosis Filtering and Identification from Live-cell Tracking data), a DL-based classifier for filtering SPT tracks of exocytic events in yeast cells expressing a subunit of the exocyst tagged with a pH-independent fluorophore. To systematically distinguish bona fide events from ambiguous events, we leveraged the learning capabilities of convolutional neural networks (CNN). We demonstrated that a CNN trained on simulated data and fine-tuned on a small experimental dataset via transfer learning achieves performance comparable to human annotators. This performance enables an overall ten-fold reduction in manual annotation effort while minimizing annotator-dependent bias. We also designed a set of user-friendly ImageJ scripts and graphical user interfaces (GUIs) to simplify the entire pipeline and to facilitate comparative analysis by other researchers. We applied ExoFILT to study the temporal relationship between the exocyst and the SM protein Sec1. Comparative analysis of exocytic events with distinctive molecular composition suggested that the number of exocysts present in each exocytic event determines the clustering of Sec1.

## 2. Methods

### 2.1. Imaging and single-particle tracking (SPT)

The experimental data used to develop ExoFILT was acquired by simultaneous dual-color live-cell imaging, as previously explained^8^. Briefly, *Saccharomyces cerevisiae* cells were imaged at the equatorial plane, expressing the exocyst subunit Exo84 tagged with three mCherry in tandem (exocyst-mCh). For this dataset, only exocyst-Ch data was considered. Time-lapse movies were acquired at 115 ms/frame for 70 seconds (615 frames) with an exposure time of 80 ms by an sCMOS Zyla 4.2 camera (Andor, Oxford Instruments) with a pixel binning set to 2 (Supplementary Note 1).

A custom pipeline was developed in ImageJ comprising (i) the preprocessing of SPT datasets and (ii) the automated tracking of exocytic events based on TrackMate^39,40^ (Supplementary Note 2). We applied this pipeline to two sets of four movies each, including approximately 350 and 370 cells per dataset (as automatically estimated after cell segmentation by StarDist^41^), generating two independent datasets of 2543 and 2458 exocyst-mCh tracks, hereafter referred to as raw datasets 1 and 2 (RD_1_ and RD_2_), respectively (Table 1). The RD_1_ dataset was manually curated by a single annotator (Ann1) using a dedicated GUI (see section 2.3), resulting in 91 tracks (3.6%) labeled as bona fide events (Table 1).

**Table 1.**
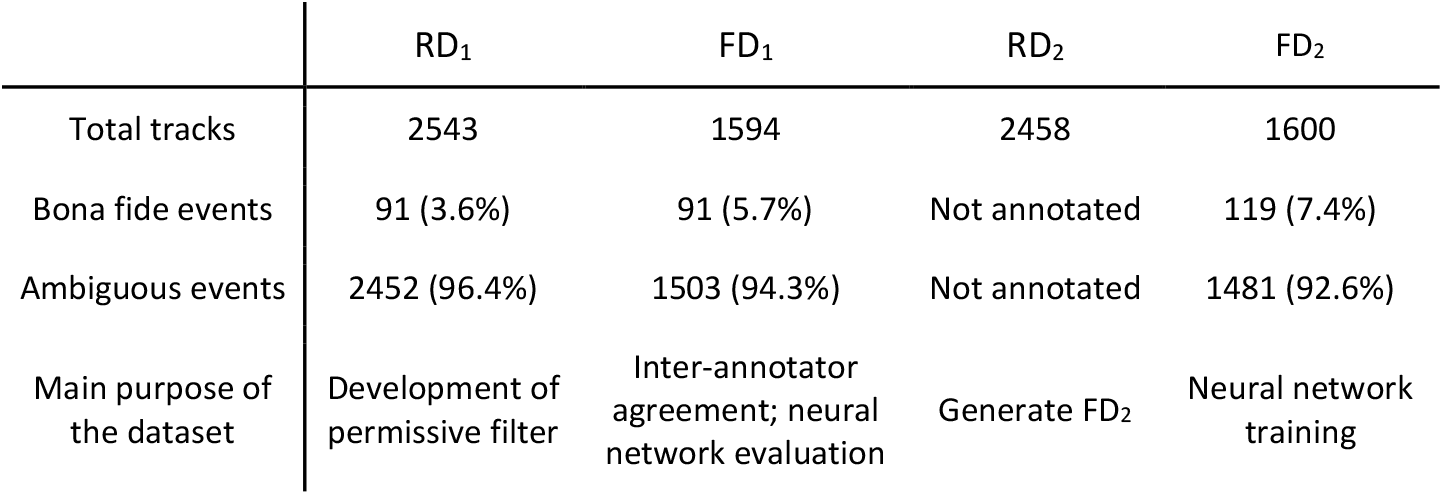
Summary of SPT datasets. Exocyst-mCh datasets annotated by Ann1, used for different purposes throughout this study. FD_1_ results from applying the permissive filter to RD_1_, while FD_2_ was annotated after applying the permissive filter to RD_2_.

For the case study, we acquired a second dataset from cells expressing exocyst-mCh together with mNeonGreen-Sec1 (mNG-Sec1) and imaged under the same conditions. The same preprocessing was conducted on the exocyst-mCh channel, in this case on 53 movies corresponding to approximately 2468 cells, yielding 79074 exocyst-mCh tracks.

### 2.2. Permissive filter

To reduce the burden of manual annotation of SPT datasets, we established a pre-selection step to exclude clearly ambiguous or spurious tracks before curation. Initial attempts relied on fixed, conservative thresholds based on quantitative track properties. These criteria were effective at removing obvious false positives but also discarded a great number of bona fide exocytic events.

To avoid losing bona fide events, we used an annotated dataset (RD_1_) to derive a permissive filter that requires no conservative thresholds but instead follows a data-driven approach (Supplementary Note 3). The permissive filter was designed to preserve nearly all bona fide events while still decreasing the overall number of tracks requiring manual annotation. The resulting permissive filter was then applied to both raw SPT datasets (RD_1_ and RD_2_), generating the filtered datasets FD_1_ and FD_2_ (Table 1).

### 2.3. GUI for manual annotation

For the quantitative analysis of exocytic events only tracks with specific characteristics were manually selected. We defined a bona fide exocytic event as a transient, nearly static exocyst-mCh diffraction-limited punctum, located near the plasma membrane, reaching a detectable fluorescence intensity above the local background, without overlaps with other events throughout its whole lifetime, and with a distinguishable start and end. Any track lacking any of these characteristics (hereafter referred to as an ambiguous event) was discarded (Fig. 1).

**Figure 1.**
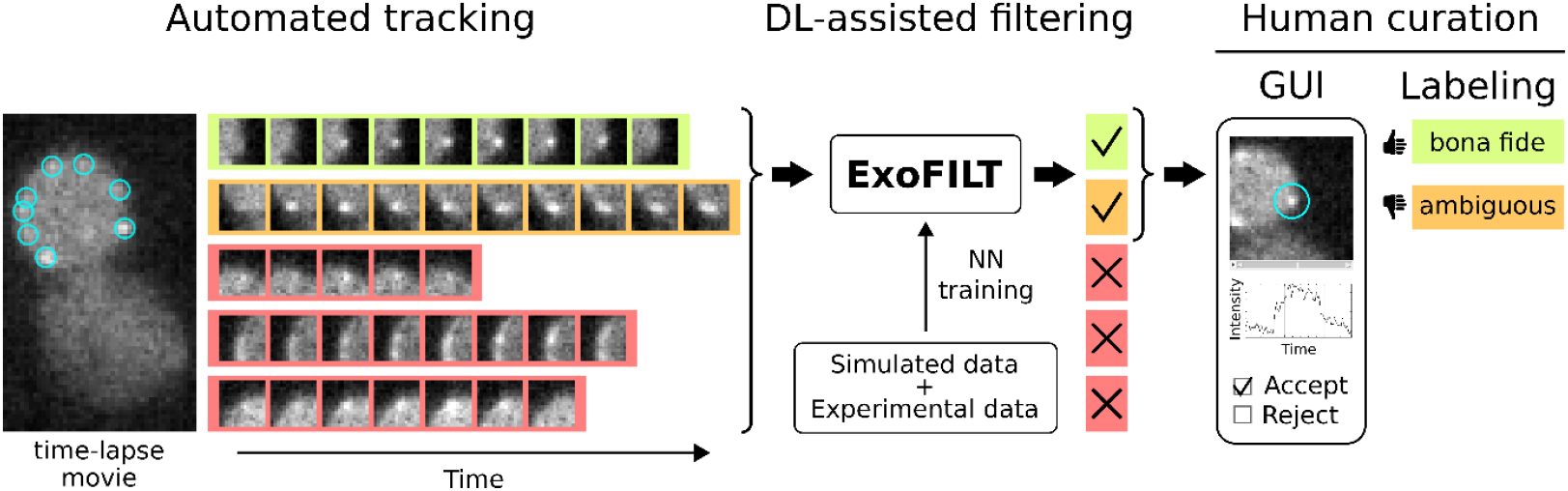
Proposed workflow for annotating exocytic events from yeast time-lapse movies. Candidate events are detected using TrackMate on preprocessed whole field-of-view time-lapse movies, each containing multiple cells. For each candidate, cropped image sequences (10×10-pixels frames) centered on the candidate event’s centroid are extracted and used as input to ExoFILT classifier, which rejects (red) or selects (green and orange) events. A user-friendly ImageJ GUI aids further curation by a human annotator, generating the final labels for each exocytic event (bona fide or ambiguous). In this example, ExoFILT correctly discarded 3 tracks, reducing the need for human curation.

Based on these qualitative criteria, exocyst-mCh tracks were manually annotated using a custom, user-friendly ImageJ-based GUI (Annotation GUI) (Fig. 1). This GUI takes as input a CSV file containing the coordinates and frame information (i.e., track start and stop as defined by TrackMate) of each track to be annotated, together with the preprocessed exocyst-mCh movies where the tracks were detected (Supplementary Note 2). After initial parameter selection, the GUI loads the list of tracks, locates each one in the corresponding movie, and displays them sequentially as cropped regions centered on the detected puncta. The track intensity profile is also extracted and displayed. The user can then annotate the event as either bona fide or ambiguous directly through the interface. By centralizing visualization and annotation within the same GUI, this tool substantially reduced the time required to curate large datasets.

In addition, we developed a complementary ImageJ GUI (Colocalization GUI) for the qualitative analysis of colocalization between channels 1 and 2, and for tracking puncta detected in channel 2. The GUI loads the CSV file of manually annotated bona fide events from channel 1 (in this study, always exocyst-mCh) and displays the corresponding cropped region for both channels side-by-side. The user can visually assess whether a bona fide event is detected in channel 2 and colocalizes with the previously annotated channel 1 event. When colocalization is observed, TrackMate processes the second channel, and a new CSV file is generated containing all detected pairs of colocalizing bona fide events. In this study, we used the Colocalization GUI to analyze a dual-color live-cell imaging dataset tracking both exocyst-mCh and mNG-Sec1.

### 2.4. Annotation Effort

We computed a metric which we named annotation effort, defined as the time required to reach 100 annotated bona fide events from a given dataset, given an average annotation speed of 5 raw tracks per minute. This annotation speed was estimated from the performance of trained researchers in our laboratory. Note that this metric is dependent only on the fraction of bona fide events in the dataset.

### 2.5. Neural Network architecture

We developed a CNN-based model aimed at improving the discrimination of bona fide exocytosis events. 3D (2D+T) convolutions were used to process short image sequences. The network was designed to handle input videos of variable length and a frame size of 10×10 pixels to preserve sufficient local context around the diffraction-limited spots. While each sample within a batch shares the same temporal length, the number of frames can vary across batches, eliminating the need for temporal cropping or padding.

The first part of the architecture is based on an Inception module^42^, with three 3D convolutional branches with kernel sizes 1x1x1, 3x3x3, and 5x5x5, applied directly to the input, together with a max-pooling branch (1x1x1 kernel with stride 1). The resulting feature maps from all branches were concatenated, followed by batch normalization and a 3D global max-pooling layer. Finally, four fully-connected sequential layers perform binary classification. A dropout layer (p = 0.5) was added after the first dense layer to reduce overfitting. See Supplementary Fig. 1 for a diagram of the architecture. Supplementary Table 1 summarizes the number of filters of the convolutional layers and the number of units of the dense layers. The model was trained using the binary cross-entropy loss function and the Adagrad optimizer. A learning rate scheduler was employed to reduce the learning rate by a factor of 0.5 when the validation loss failed to improve after 20 epochs.

### 2.6. Neural Network training

All models were trained on a workstation equipped with an NVIDIA Quadro RTX 4000 GPU and an Intel Xeon Gold 6238R CPU. The framework was implemented in Python 3.9 using Keras (v2.10.0)^43^ with a TensorFlow backend^44^ and Weights & Biases^45^ was used for experiment tracking and visualization. We tested the same neural network model under three training strategies: (A) training exclusively on experimental data, (B) training exclusively on simulated data (i.e., synthetically generated exocytic events, see Section 2.8), and (C) a transfer learning strategy (ExoFILT).

To train the previously described model on experimental data (FD_2_), either directly (training strategy A) or via transfer learning (ExoFILT), we performed five runs using 5-fold cross-validation with an 80:20 train-validation split. To increase robustness and compensate for the limited amount of experimental data, we applied data augmentation by generating six variants of each input video through 90° rotations and mirroring along two perpendicular axes. During training, these augmented videos were treated as independent samples. For inference, the same augmentation scheme was applied to each input video, and predictions across the six variants were averaged into a single score per training run. The final score of each training modality was computed as the mean score across all five runs. When training the network only on simulated data (training strategy B), no cross-validation was used but five identical runs were performed. In all training modalities, no model among the five training runs was found to be better or worse than the others.

For training strategy A, a batch size of 6 was used. The initial learning rate was set to 0.01. The training dataset was class-balanced, as an imbalanced sampling (1 bona fide: 10 ambiguous videos) led to a collapse to the majority class. In total, 112 videos from each class were used for training and validation. The training size was limited by the bona fide events available in FD_2_ (Table 1).

For training strategy B, the batch size was increased to 64 and the learning rate was set to 0.001. We evaluated eight different training set sizes ranging from 1000 to 150000 samples (always using balanced class sampling). Classification accuracy on the test set increased with training set size up to approximately 50000 samples, after which performance plateaued, while training time continued to increase. Based on this observation, a training size of 50000 was selected. This required a simulated dataset of 71430 videos, which was split into training (70%), validation (15%), and test (15%) sets.

For the transfer learning approach (ExoFILT), the batch size was set to 6, as in strategy A, while the learning rate was reduced to 0.0005. Instead of random weight initialization, one of the five models trained under strategy B (simulated data) was used as the starting point for the training. All trainable layers (i.e., dense and convolutional layers) were unfrozen, as this yielded slightly better results than partial freezing of the network. The network was fine-tuned using experimental data from dataset FD_2_. For this scenario, an imbalanced class sampling approach (1 bona fide: 10 ambiguous videos) was used as it led to a slight performance improvement over a balanced sampling approach (Supplementary Fig. 2). For training and validation, the total number of bona fide and ambiguous videos was respectively 112 and 1125.

### 2.7. Performance metrics

Here we report the formulas for the metrics used throughout this work to evaluate inter-annotator agreement and model performance. The metrics are defined as follows, where TP, FP, TN and FN are the number of true positives, false positives, true negatives, and false negatives, respectively (positive = bona fide; negative = ambiguous):

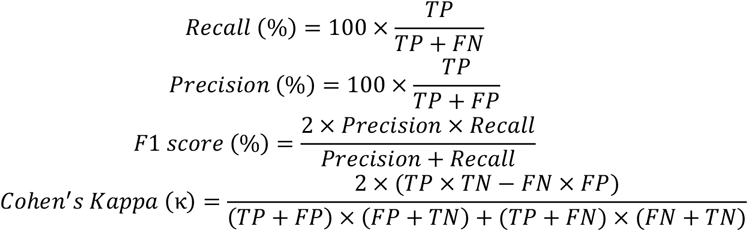

Recall measures the fraction of bona fide events correctly detected, whereas precision quantifies the fraction of predicted bona fide events that are correct. F1 score provides a single summary metric that balances these two quantities and is particularly useful under class imbalance. Cohen’s Kappa (κ) assesses agreement beyond chance, allowing comparisons between annotators or between model predictions and ground truth.

To evaluate performance across thresholds, we also report the area under the receiver operating characteristic curve (AUC-ROC), which summarizes the trade-off between true positive rate and false positive rate, and the area under the precision-recall curve (AUC-PR), which is especially informative in imbalanced scenarios^46^.

### 2.8. Data simulations

We simulated movies mimicking experimental exocytic events, both bona fide and ambiguous (Supplementary Fig. 3). Simulations were based on DeepTrack 2.1^47^ and the AnDi-Datasets Python package^48^. Sequences of 10×10-pixels frames were obtained by varying a wide set of parameters. Further details on key aspects of these simulations are described in Supplementary Note 4. By combining a variety of features from the described pipeline, we defined 3 subclasses of simulated bona fide exocytic events and 11 subclasses of ambiguous events (Supplementary Table 2). We generated 20000 videos for each subclass, which provided sufficient simulated data to train our model (training strategy B).

**Table 2.**
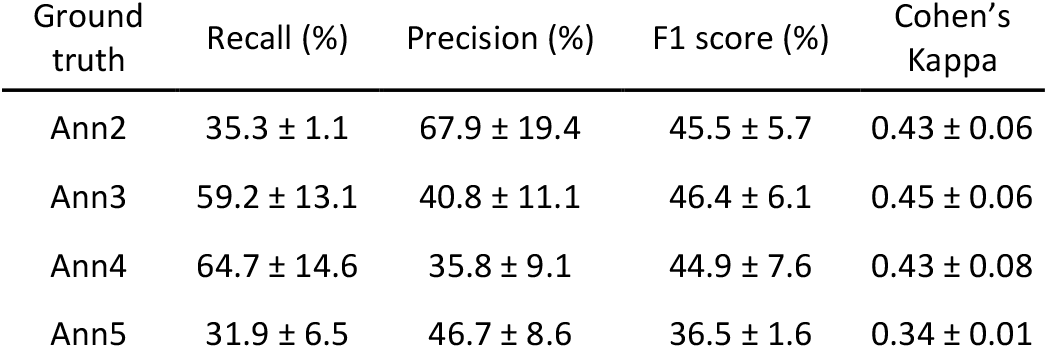
Inter-annotator agreement. Average human-human recall, precision, F1 score, and Cohen’s Kappa (κ). For each ground truth, we report the mean ± SD from the pairwise comparisons with the three other annotators (see Supplementary Table 3 for raw pairwise comparisons).

## 3. Results

### 3.1. Construction of SPT datasets

To image exocytosis, we used the exocyst as a temporal reference due to its central role within the pathway, from initiation of the tethering to vesicle fusion^8,9^. *S. cerevisiae* cells expressing the exocyst fused to three mCherry in tandem (exocyst-mCh) were imaged (Fig. 1 and section 2.1). Upon SPT analysis, two raw datasets were generated (RD_1_ and RD_2_).

We applied the permissive filter (section 2.2, Supplementary Note 3) to RD_1_, resulting in a filtered dataset (FD_1_) of 1594 tracks (91 bona fide exocytic events and 1503 ambiguous, as annotated by Ann1), where 38.7% of the ambiguous tracks in RD_1_ were effectively removed (Table 1). FD_1_ was used for further analyses, including inter-annotator agreement assessment and evaluating a DL-based classifier. We then applied the permissive filter on RD_2_, resulting in a filtered dataset (FD_2_) of 1600 tracks (119 bona fide exocytic events and 1481 ambiguous, as annotated by Ann1). This set was used for training our DL classifier.

### 3.2. Inter-annotator agreement, biases and annotation effort

Using FD_1_, we estimated the inherent variability of human annotation. As mentioned above, both technical and biological limitations make manual annotation both intensive and subjective. To estimate this variability, four researchers (Ann2-Ann5) independently annotated the same dataset. The average annotation effort (section 2.4) was 10.7 ± 4.5 h (Fig. 2A). Pairwise metrics (recall, precision, F1 score, and Cohen’s Kappa κ) were computed by treating one annotator as the reference (ground truth) and the other as the prediction.

**Figure 2.**
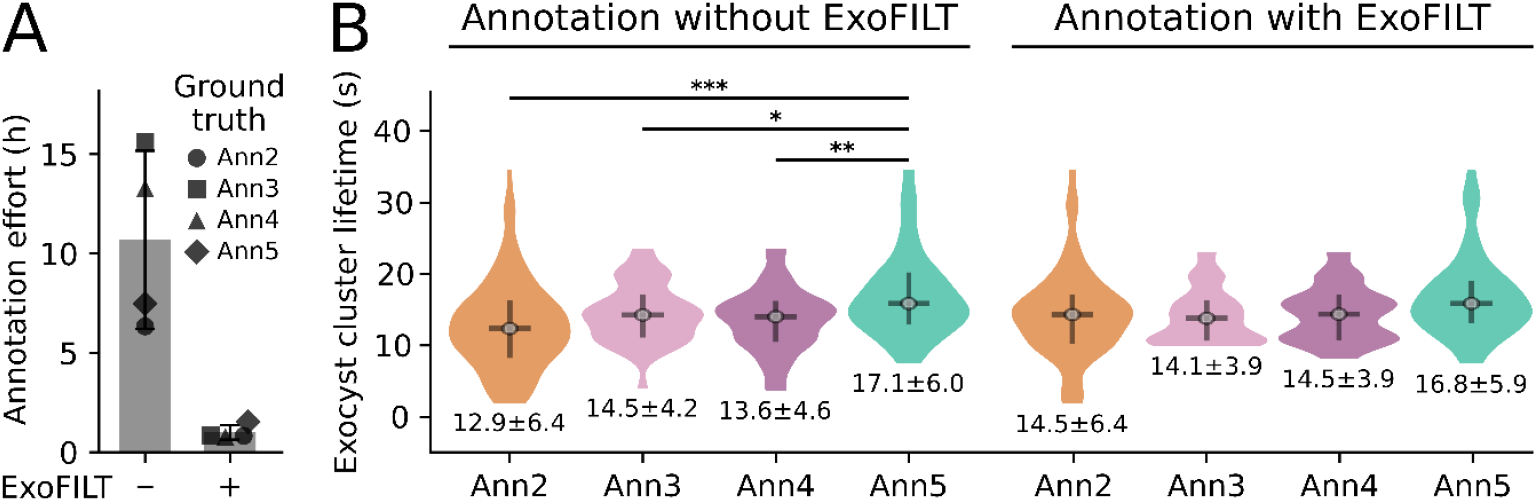
Annotation of exocytic events before and after applying ExoFILT. (A) Annotation effort on dataset FD_1_, using either the permissive filter alone or combined with ExoFILT. (B) Violin plots of exocyst cluster lifetime for bona fide events selected by indicated annotators, before (left) and after (right) applying ExoFILT. The corresponding mean ± SD exocyst cluster lifetime is shown below each distribution. The horizontal bar indicates the median, and the vertical bar corresponds to the interquartile range (25th-75th percentiles). Number of events by annotator are summarized in Supplementary Table 4. Statistical significance (Mann-Whitney U test, two-sided) is indicated as ***p<0.001, **p<0.01, *p<0.05 (non-significant comparisons are not displayed).

Overall, 132 tracks were annotated as bona fide by at least one annotator, but only 12 were labeled as bona fide by all annotators, reflecting strong personal biases. Consequently, averaging across the six pairwise comparisons obtained by switching the ground truth annotations yielded a mean F1 score of 43.3 ± 6.9%, while κ averaged 0.41 ± 0.07 (Table 2).

To better understand the origin of human biases, we examined recall and precision of each researcher as an evaluator across all pairwise comparisons (Supplementary Table 3). This revealed distinct annotation strategies: Ann2 consistently showed the highest recall for each ground truth, indicative of a more permissive interpretation of bona fide exocytic events, whereas Ann4 exhibited the highest precision, reflecting a more conservative approach.

Importantly, despite the low reproducibility of the annotations, no significant difference could be detected for the average lifetime of bona fide exocyst-mCh puncta selected by Ann2 (12.9 ± 6.4 s), Ann3 (14.5 ± 4.2 s) and Ann4 (13.6 ± 4.6 s). Consistent with a significant bias introduced by Ann5, the average lifetime of bona fide exocyst-mCh puncta derived from this annotator (17.1 ± 6.0 s) showed a significant deviation (Fig. 2B), with an average F1 score of 36.5 ± 1.6% and κ of 0.34 ± 0.01 when used as ground truth (Table 2). These differences allowed us to establish a realistic human-level baseline to contextualize the performance of a DL-based classifier. Overall, this comparative analysis illustrates the complexity of annotating this type of data, even for expert annotators, and the risk of introducing personal biases that impact the measurement of biological features, restricting its suitability for systematic and quantitative analyses of SPT of exocytosis.

### 3.3. DL-based classifier to filter exocytosis SPT datasets

We developed a DL-based binary classifier following a CNN architecture, using 3D (2D+T) convolutions to learn both spatial and temporal features from the data. The model labeled tracks as either bona fide or ambiguous events (section 2.5, Supplementary Fig. 1).

We set the model input spatial dimensions to 10×10 pixels windows cropped around the track centroid. This simplified the pipeline for generating simulated training data, but at the cost of limiting network depth. We addressed this by incorporating an Inception module^42^, which applies multiple convolutional operations with different kernel sizes in parallel. After concatenating the branches from the Inception module, the subsequent layers followed a classical structure consisting of batch normalization, 3D global max-pooling, and dense layers for binary classification, with a dropout layer after the first dense layer to reduce overfitting (section 2.5, Supplementary Fig. 1).

### 3.4. Neural Network training strategies and performance evaluation

To train the CNN-based classifier we followed three strategies: (A) training exclusively on experimental data (FD_2_), (B) training exclusively on simulated data, and (C) a transfer learning strategy (ExoFILT) in which a model trained on simulated data was fine-tuned using experimental data.

We used an independent experimental dataset (FD_1_) to test performance across strategies and evaluate their generalization capability. Using each of Ann2-Ann5 annotations as ground truth, we evaluated each training strategy with AUC-ROC and AUC-PR, and a decision threshold maximizing F1 score (optimal decision threshold) was chosen to compare both F1 score and κ across strategies and against human performance.

#### 3.4.1. Training strategy A: Training on experimental data

We first assessed whether the available experimental data was sufficient to train our neural network model. For this purpose, we used FD_2_, annotated by Ann1, for training and validation. We trained our models on a balanced dataset (section 2.6), reaching an average accuracy of 93.5 ± 1.4% on the training set and 75.2 ± 4.0% on the validation set. This difference, together with the learning curves (Supplementary Fig. 4), suggests that the model might be overfitting.

On FD_1_, AUC-ROC was 0.86 ± 0.03, while AUC-PR was substantially lower (0.26 ± 0.09) (Fig. 3A,B; Supplementary Fig. 5). This difference highlights the relevance of using both metrics for imbalanced classification tasks, as AUC-ROC can remain high even when precision or recall are low. At the optimal decision threshold, the average F1 score and κ were 32.1 ± 6.2% (Fig. 3C) and 0.29 ± 0.08 (Supplementary Fig. 6), respectively. Overall, these results suggest that the amount of available experimental data is insufficient to train the model to generalize and reach human-level performance.

**Figure 3.**
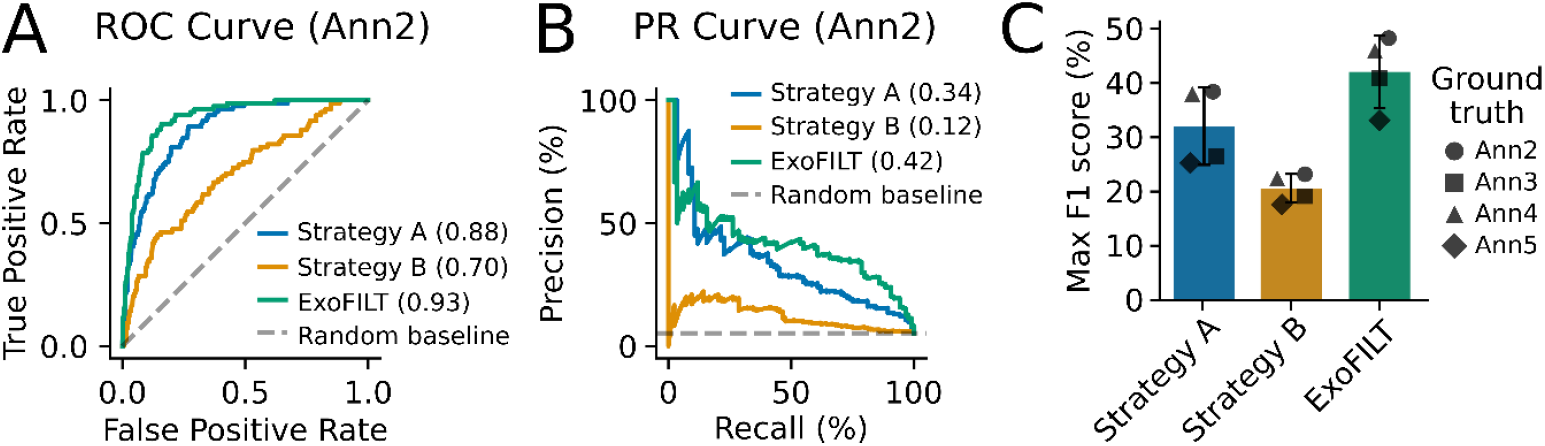
Performance metrics for different training strategies. ROC curve (A) and PR curve (B) for the three training strategies, given a representative ground truth (Ann2). See Supplementary Fig. 5 for the rest of annotators. AUC values for each strategy are shown in the legend. The random baseline in the PR curve corresponds to the prevalence of the bona fide class (i.e., percentage of events annotated as bona fide by Ann2, which is 0.05). (C) Maximum F1 score per strategy for each ground truth annotator. The height of the bar represents the mean across ground truths, while the error bars represent the standard deviation.

#### 3.4.2. Training strategy B: Training on simulated data

We then tested whether simulated time-lapse videos of exocytic events could be employed to directly train our model (Supplementary Fig. 3, Supplementary Note 4). We trained the neural network on a balanced simulated dataset of 50000 samples, achieving an average accuracy of 98.0 ± 0.2% on both training and validation sets (Supplementary Fig. 4). These results emphasize the models’ ability to accurately learn from large datasets, but might also reflect the internal consistency and relative simplicity of the simulated data.

On FD_1_, all performance metrics were lower than those obtained with strategy A. AUC-ROC and AUC-PR were 0.71 ± 0.06 and 0.10 ± 0.01, respectively (Fig. 3A,B; Supplementary Fig. 5). The maximum F1 score was 20.6 ± 2.3% (Fig. 3C), with a corresponding κ of 0.17 ± 0.02 (Supplementary Fig. 6). Despite the low performance, the stability of the model trained on extensive simulated data suggested that these models could serve as a starting point for a transfer learning task.

#### 3.4.3. Training strategy C (ExoFILT): A transfer learning strategy

We used a single model trained on simulated data (strategy B) as a base model and fine-tuned it with experimental data (FD_2_), following a transfer learning approach (section 2.6). This technique has been widely applied in bioimage analysis when annotated data is limited^49–53^. The resulting model was named ExoFILT (Exocytosis Filtering and Identification from Live-cell Tracking data).

To address the class imbalance between bona fide and ambiguous events in the experimental data, we compared two sampling approaches: balanced sampling (1:1 class ratio) and imbalanced sampling (1:10). When training directly on experimental data (strategy A), imbalanced sampling led to a collapse to the majority class. In contrast, under the transfer learning scenario, imbalanced sampling was stable and resulted in a modest performance improvement compared with balanced sampling (Supplementary Fig. 2). We therefore used a 1:10 sampling ratio to train ExoFILT, as it reflects better the class distribution observed in experimental data. Although stronger effects might be achieved under more extreme imbalance ratios, the available training data did not allow us to systematically evaluate them.

With this configuration, training was stable and consistent across cross-validation runs, with average training and validation accuracies of 94.2 ± 0.4% and 93.1 ± 0.7%, respectively, with no signs of overfitting (Supplementary Fig. 4). Performance on FD_1_ also improved substantially. AUC-ROC and AUC-PR reached 0.92 ± 0.02 and 0.33 ± 0.07, respectively (Fig. 3A,B, Supplementary Fig. 5). The optimal decision threshold yielded an average F1 score of 42.1 ± 5.8% (Fig. 3C) and κ of 0.39 ± 0.07 (Supplementary Fig. 6). Overall, ExoFILT’s agreement with individual annotators was comparable to inter-annotator agreement (Table 2).

We examined ExoFILT’s robustness to the decision threshold by measuring recall, precision and F1 score across thresholds between 0 and 1. F1 score stayed within 25% of its maximum for a wide range of thresholds (average across annotators: 38.2 ± 11.2% of all possible thresholds) (Supplementary Fig. 7A), indicating robustness to threshold selection. This robustness gives flexibility to the user to adapt the threshold to favor either precision or recall, useful in scenarios where annotation time is limited (favoring precision) or where there is a lack of sufficient bona fide events (favoring recall).

The assistance of ExoFILT reduced about 10-fold the annotation effort for FD_1_ across annotators (1.0 ± 0.4 hours; Fig. 2A). With ExoFILT, no significant differences in exocyst-mCh lifetimes across annotators could be detected (Fig. 2B, right). These results indicate that ExoFILT can help mitigate personal biases and therefore facilitate comparative analysis of exocyst-mCh puncta lifetime measurements.

### 3.5. Case study: Quantitative dynamics of exocyst and Sec1 illuminate a mechanism controlling exocytosis functionality

ExoFILT opens new opportunities to investigate the temporal relationship between the exocyst and other components of the exocytic machinery. Here, we applied ExoFILT to simultaneous dual-color time-lapse movies of cells expressing exocyst-mCh and Sec1 tagged with mNeonGreen (mNG-Sec1). By comparing distinct populations of exocytic events, we aimed to identify molecular features that contribute to exocytosis functionality.

Following SPT analysis on a dataset of 53 movies (section 2.1), we applied the permissive filter to the exocyst-mCh channel, yielding a filtered dataset of 21002 tracks. These were subsequently processed with ExoFILT using a decision threshold of 0.35, selected to balance recall and annotation effort. This resulted in 412 candidate events selected for manual inspection. Using the Annotation GUI (section 2.3), a single annotator (Ann1) identified 194 exocyst-mCh bona fide events (annotation effort of 0.7 hours).

Colocalization with mNG-Sec1 was then assessed using the Colocalization GUI (section 2.3). Among the 194 exocyst-mCh bona fide events, 98 colocalized with a mNG-Sec1 bona fide event, defined using the same criteria as for exocyst-mCh. Of the remaining exocyst-mCh events, 52 colocalized with an ambiguous mNG-Sec1 track and 44 showed no detectable colocalization with mNG-Sec1 signal. Thus, 22.7% of bona fide exocyst-mCh events lack detectable Sec1 recruitment. This is in agreement with previous studies indicating that a substantial fraction of exocytic events do not culminate with vesicle fusion (i.e. they are abortive events) or might follow alternative mechanisms^8,24^. Such heterogeneity of exocytic events has been overlooked in the past due to the lack of data.

Using the 98 bona fide exocyst-mCh/mNG-Sec1 pairs, we computed median intensity profiles as well as clustering and disassembly times for puncta of both proteins (Fig. 4A,B and Supplementary Note 5). Under our experimental conditions, exocyst-mCh events have an average lifetime of 13.4 ± 6.6 s, consistent with the dataset used to assess inter-annotator agreement (Fig. 2B). We observed that mNG-Sec1 puncta, with an average lifetime of 8.8 ± 4.8 s, are recruited at exocytic events 5.1 ± 5.4 s after exocyst-mCh clustering. The mNG-Sec1 intensity profile displays a peak during the final ∼2 s of the exocyst-mCh lifetime. Finally, the mNG-Sec1 puncta dissociates simultaneously to the exocyst-mCh disassembly.

**Figure 4.**
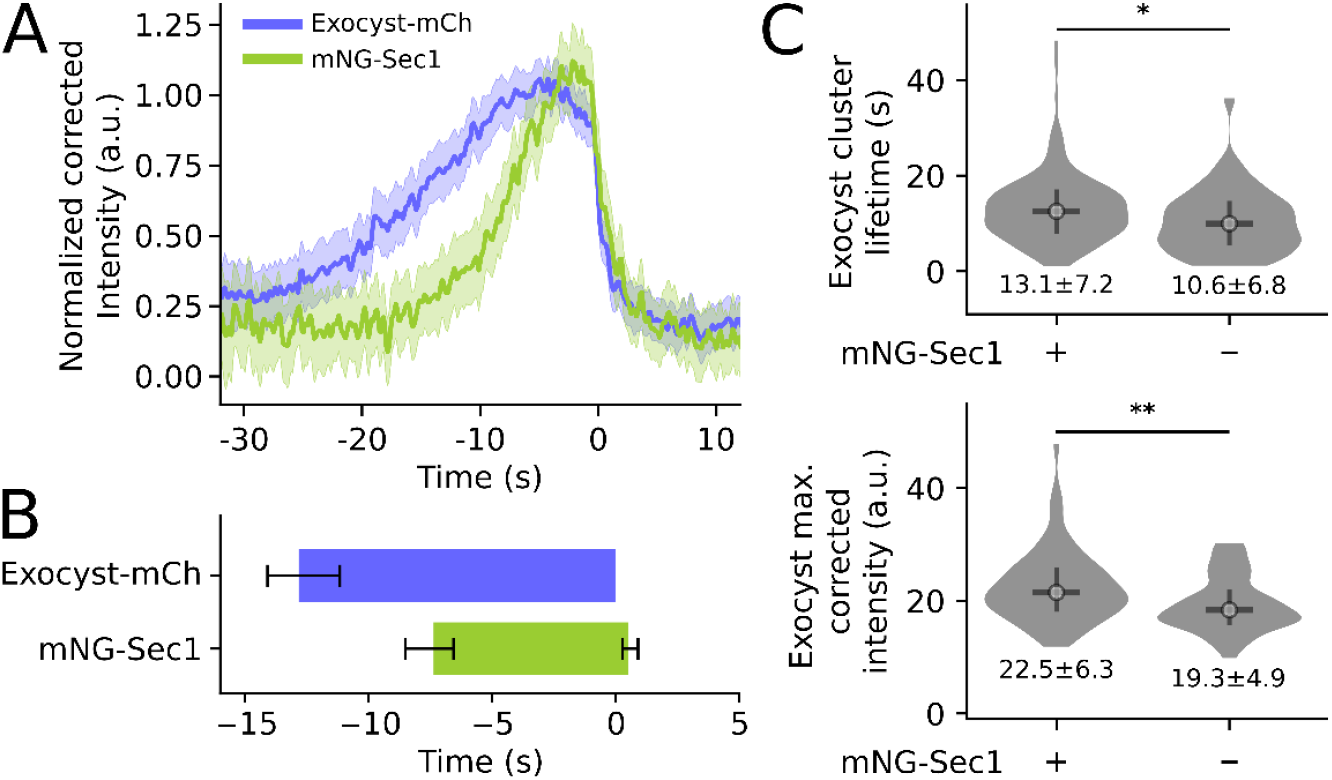
Quantitative timeline of the exocyst and Sec1. (A) Average intensity profiles and (B) timelines for colocalizing bona fide exocyst-mCh and mNG-Sec1 event pairs (n = 98). In panel A, the median intensity at each timepoint is shown; shaded area indicates the 95% confidence interval. Panel B displays the median clustering and disassembly times; error bars represent the 95% confidence interval. (C) Comparison of cluster lifetime (upper panel) and maximum corrected intensity (lower panel) of exocyst-mCh events colocalizing with bona fide or ambiguous mNG-Sec1 events (n = 150) and not colocalizing with detectable mNG-Sec1 (n = 44). Values shown below each distribution indicate mean ± SD. The central horizontal bar indicates the median, and the vertical bar corresponds to the interquartile range (25th-75th percentiles). Statistical significance (Mann-Whitney U test, two-sided) is indicated as **p<0.01 and *p<0.05.

We next compared exocyst-mCh events that colocalized with mNG-Sec1 (including bona fide and ambiguous mNG-Sec1 events) to those without detectable mNG-Sec1 signal (Supplementary Note 5). mNG-Sec1-negative events exhibit significantly shorter lifetimes (10.6 ± 6.8 s) than mNG-Sec1-positive events (13.1 ± 7.2 s) (Fig. 4C, upper panel). In addition, their peak exocyst-mCh intensity, a proxy for the maximum copy number of exocyst molecules per event, is also significantly lower (Fig. 4C, lower panel). These results suggest that mNG-Sec1-negative events represent a distinct class of exocyst clusters that do not correspond to canonical exocytic events. Clusters harboring fewer exocyst copies are less likely to recruit mNG-Sec1 and are rapidly disassembled, likely failing to complete vesicle fusion. This interpretation is consistent with previous work showing that the levels of cytosolic exocyst available to undergo clustering limits the rate of initiation of new exocytic events^8^.

## 4. Discussion

ExoFILT is a transfer learning-based framework designed to simplify and accelerate the annotation of exocytosis events in yeast cells, thus improving the robustness and throughput of SPT-based analysis. By integrating ExoFILT with a set of user-friendly ImageJ scripts and GUIs for preprocessing and annotation, we provide a practical and accessible pipeline for the quantitative analysis of exocytosis (Fig. 1).

We trained ExoFILT on exocyst-mCh SPT data because this protein complex presents unique properties that multiply the applicability of this tool: 1) The exocyst is conserved and strictly necessary for the initial tethering of secretory vesicles to the plasma membrane. Thus, by tracking exocyst-mCh diffraction-limited puncta we can cover a large space of physiologically relevant scenarios, even when the secretory vesicle is not fused. 2) Its relatively long lifetime at the exocytic event aids the reliable assignment to this same event of the protein imaged in the second channel. 3) Exocyst clusters associated to the plasma membrane have been structurally resolved^8,19,24^, including their dynamics and stoichiometry, which makes the exocyst-mCh diffraction-limited puncta a quantifiable standard reference for both yeast and mammalian cells. Overall, we believe that the exocyst tagged with a pH-independent fluorophore constitutes a valuable reference system to understand the temporal relationships and dynamics of the exocytic machinery.

ExoFILT opens the possibility of investigating the physiological heterogeneity of different exocytic events, a mechanistic insight that had been greatly overlooked in the past. Although it was only trained and validated on the exocyst-mCh SPT analysis, ExoFILT can be used to derive quantitative measurements of other components of the exocytic machinery using simultaneous imaging. In this regard, the combination of ExoFILT with simultaneous dual-color imaging of exocyst-mCh and mNG-Sec1 allowed us to investigate, with greater statistical power, the molecular principles that underlie different types of exocytic events. Beyond reproducing the dynamics of mNG-Sec1^8^, we found that 22.7% of the exocytic events, under our experimental conditions, present a differential Sec1 composition (Fig. 4C). The exact fate of these exocytic events goes beyond the scope of the present study, as it would require imaging the ultrastructure of these events, a task only achievable by electron microscopy. Nonetheless, the analysis of the time-resolved intensity of these tracks allowed us to hypothesize that they are abortive exocytic events that do not complete vesicle fusion presumably because they miss enough copies of the exocyst complex. In agreement with this hypothesis, exocytic events within the lowest quartile of maximum intensity were enriched for exocyst-mCh bona fide events lacking detectable mNG-Sec1 signal (40.8% vs 16.6%; odds ratio = 3.48; two-sided Fisher’s exact test, p = 0.0013; Supplementary Note 5).

ExoFILT currently misses 44.8 ± 11.5% of exocyst-mCh bona fide events (Supplementary Fig. 7B), highlighting the need for further improvement of this tool. Reducing discrepancies between simulation and experimental data (i.e., modeling more realistic particle dynamics, heterogeneous backgrounds, or photobleaching kinetics) could enhance both recall and precision. In parallel, generative or diffusion-based models trained on annotated datasets may help synthesize rare or borderline exocytic events that are underrepresented in real data, increasing robustness without substantially increasing manual annotation effort. Finally, refinements in the model’s architecture, such as incorporating attention mechanisms to better capture spatiotemporal dynamics or integrating image restoration strategies, could further strengthen performance, also across diverse biological applications.

Exocytosis is not the only cellular pathway that occurs in diffraction-limited puncta at the plasma membrane. Endocytosis, unconventional secretion, and eisosome-associated processes represent additional contexts in which ExoFILT could be applied. Although its performance on SPT data from these pathways remains to be evaluated, adapting a model pretrained on our simulated dataset followed by fine-tuning on pathway-specific experimental data may provide a practical strategy. Minor adjustments to the simulation pipeline may further improve transferability to new datasets. We thus anticipate that the framework presented here may inspire researchers in the SPT community facing similar low-SNR challenges to adapt this strategy to their own biological systems.

With ExoFILT, we aim to make quantitative cell biology of exocytosis more accessible. Although the advancement of quantitative cell biology of exocytosis will also depend on tightly controlled experimental setups, these must be supported by robust and scalable computational analysis. By substantially reducing the annotation burden and improving throughput, robustness, and reproducibility of SPT measurements, ExoFILT alleviates a major bottleneck in exocytosis image analysis, enabling quantitative studies at a scale that had been difficult to achieve. Thus, we believe that ExoFILT will facilitate the systematic comparison of exocytosis dynamics across physiological conditions (e.g., environmental perturbations, mutants, etc), thereby expanding the range of biological insight that can be investigated.

## Acknowledgements

We thank Altair C. Hernández and Sebastian Ortiz for helpful discussions on data analysis.

## Author contributions

Conceptualization, E.K., C.M., B.O., and O.G.; methodology, E.K.; software, E.K.; data curation, E.K., L.I.B., S.M., and S.T.; formal analysis, E.K.; investigation, E.K.; resources, O.G.; writing - original draft, E.K.; writing - review & editing, E.K., L.I.B., S.T., C.M., B.O., and O.G.; supervision, C.M., B.O., and O.G.; project administration, O.G.; and funding acquisition, B.O., and O.G.

## Data availability

All raw data and code used for this manuscript will be available for reviewers during revision and when the work is published it will be available in GitHub (https://github.com/GallegoLab/ExoFILT).

