## Supplementary Material for "ExoFILT: Transfer learning for robust and accelerated analysis of exocytosis single-particle tracking data"

***Supplementary Figures***
*Supplementary Figure 1. Neural network architecture.*
*Supplementary Figure 2. Comparison of balanced versus imbalanced sampling for the transfer learning approach.*
*Supplementary Figure 3. Diagram of the data simulation pipeline.*
*Supplementary Figure 4. Learning curves for three training strategies.*
*Supplementary Figure 5. Threshold-independent metrics: ROC and PR curves.*
*Supplementary Figure 6. Cohen's Kappa coefficient (κ) for 3 training strategies across annotators.*
*Supplementary Figure 7. ExoFILT metrics across decision thresholds.*

***Supplementary Tables***
*Supplementary Table 1. Neural network architecture parameters*
*Supplementary Table 2. Subclasses of simulated exocytic events.*
*Supplementary Table 3. Inter-annotator agreement assessment*
*Supplementary Table 4. Number of bona fide events.*

***Supplementary Notes***
*Supplementary Note 1: Data acquisition*
*Supplementary Note 2: Data preprocessing and automated tracking*
*Supplementary Note 3: Permissive filter definition*
*Supplementary Note 4: Data simulations*
*Supplementary Note 5: Intensity profiles and colocalization of exocyst-mCh and mNG-Sec1*

### Supplementary Figure 1

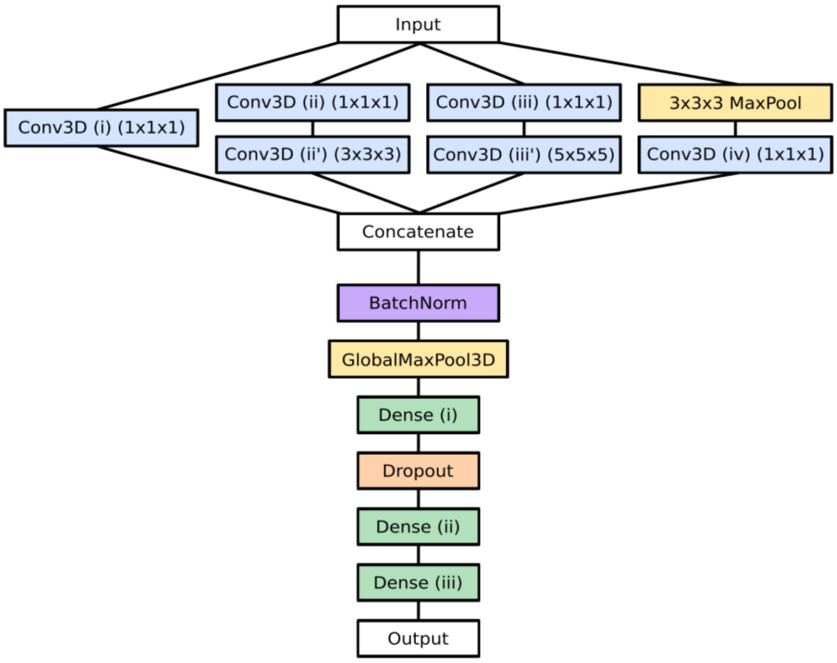


***Supplementary Figure 1. Neural network architecture.***
*Schematic representation of the 3D convolutional neural network architecture used to classify bona fide and ambiguous exocytic events from live-cell imaging data.*

### Supplementary Figure 2

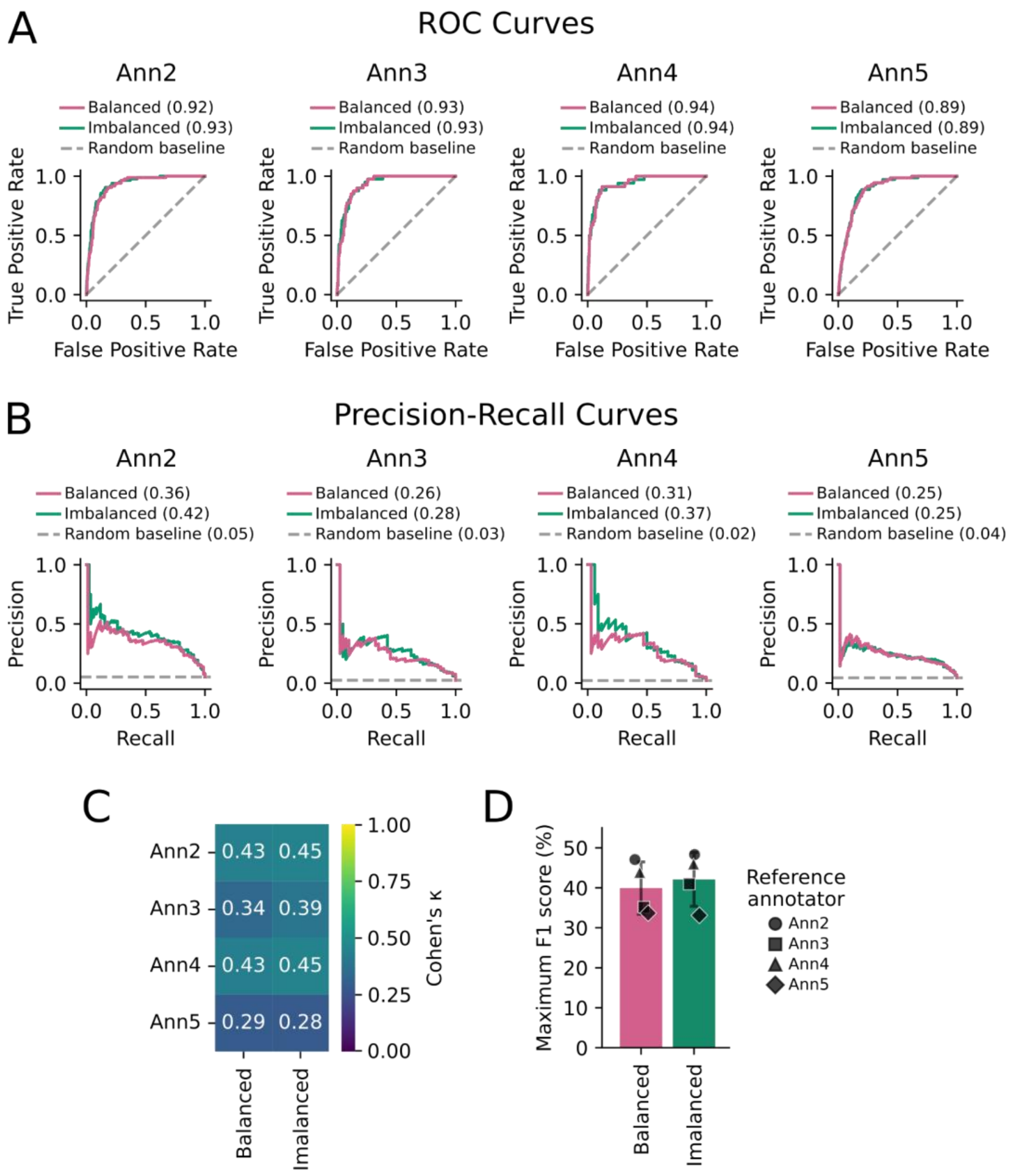


***Supplementary Figure 2. Comparison of balanced versus imbalanced sampling for the transfer learning approach.***

*ROC curves (A) and PR curves (B) for each ground truth annotator (Ann2-Ann5). For each ground truth, the optimal threshold (i.e., maximum F1 score) was selected to compute both Cohen's Kappa (C) as well as F1 score (D). Note that transfer learning with imbalanced sampling (1:10 class ratio) is the strategy we named as ExoFILT throughout the rest of the study.*

### Supplementary Figure 3

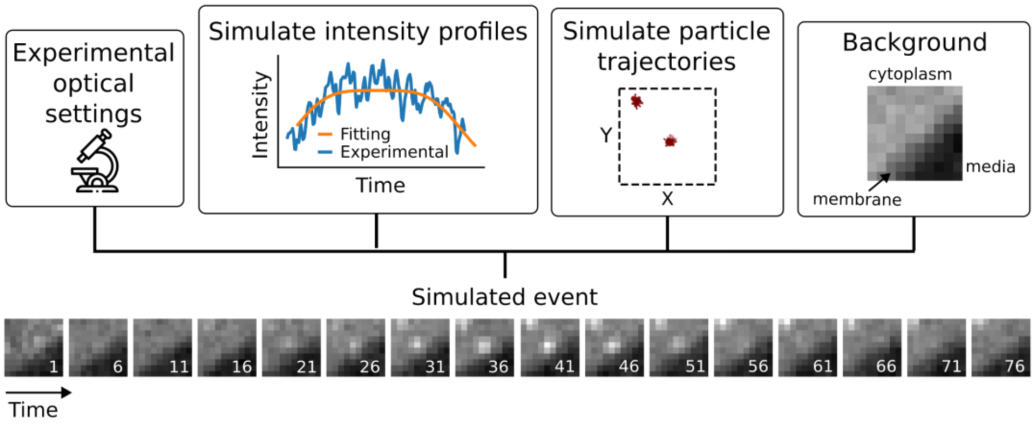


***Supplementary Figure 3. Diagram of the data simulation pipeline.***

*The integration of multiple parameters enables the generation of a heterogeneous simulated dataset that mimics exocyst-mCh tracks, both bona fide and ambiguous events. As an example, a simulated bona fide event is shown (numbers indicate the frame number within the movie).*

### Supplementary Figure 4

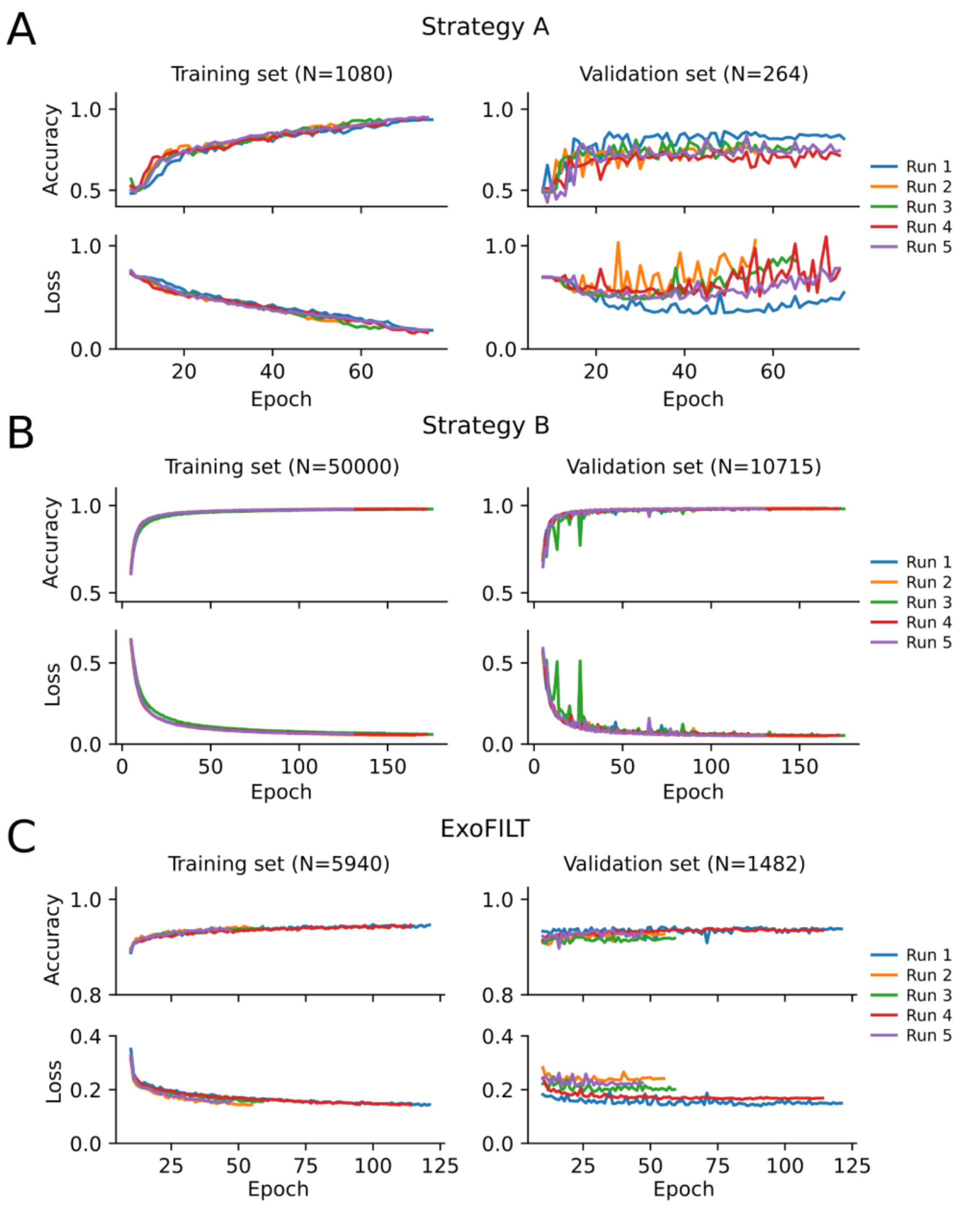


***Supplementary Figure 4. Learning curves for three training strategies.***

*Learning curves for 5 runs for the three training strategies: (A) strategy A, (B) strategy B, and (C) ExoFILT. For (A) and (C), the reported dataset sizes correspond to the number of samples after data augmentation (6 versions per original video). For (B), the reported value directly corresponds to the number of original simulated videos, since this strategy did not involve data augmentation for training or validation.*

### Supplementary Figure 5

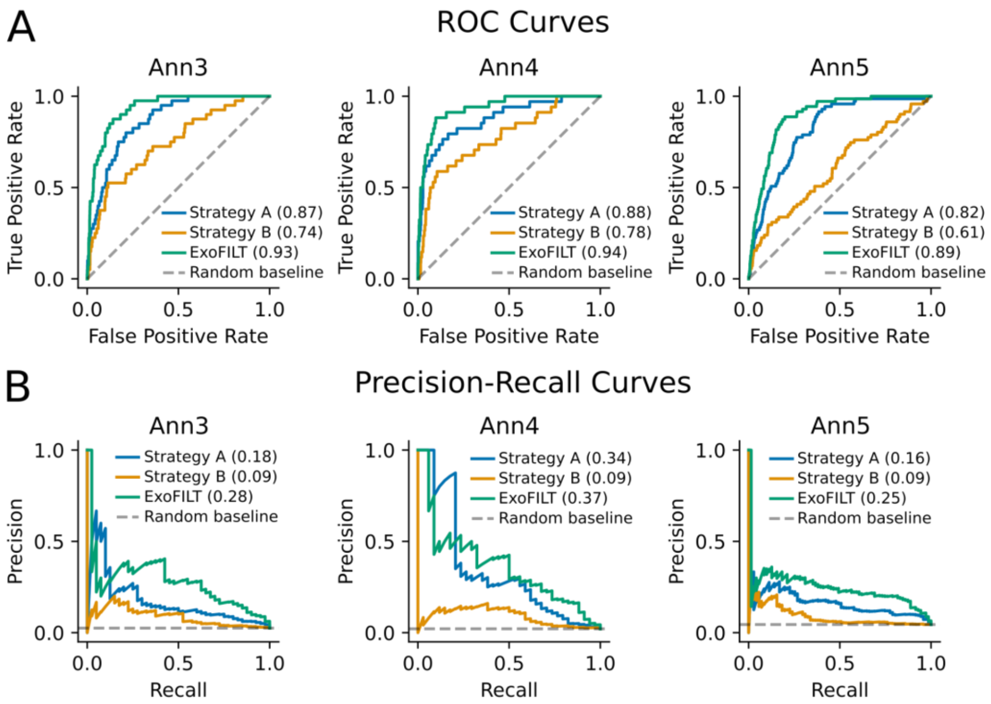


***Supplementary Figure 5. Threshold-independent metrics: ROC and PR curves.***
*Receiver-operating characteristic (ROC) curves (A) and Precision-Recall (PR) curves (B) for each training strategy evaluated against annotators Ann3, Ann4, and Ann5 (see Fig. 3 for Ann2). In the PR curves, the random baseline corresponds to the prevalence of the bona fide class (i.e., fraction of events annotated as bona fide by the corresponding annotator), which is 0.03, 0.02, and 0.04 for annotators Ann3, Ann4, and Ann5, respectively.*

### Supplementary Figure 6

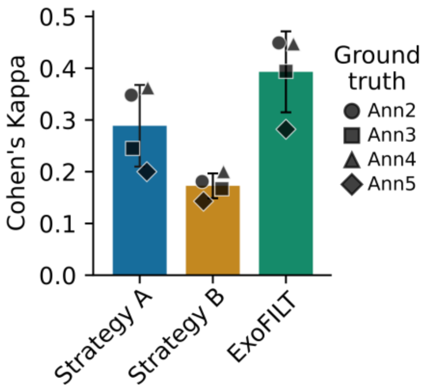


***Supplementary Figure 6. Cohen's Kappa coefficient (κ) for 3 training strategies across annotators.*** *Agreement between the neural network trained under 3 strategies and annotators Ann2-Ann5, as measured with Cohen's Kappa. This metric reflects the agreement between two sets of binary annotations, with values ranging from 1 (perfect agreement) to -1 (perfect disagreement), with 0 representing agreement expected by random chance.*

### Supplementary Figure 7

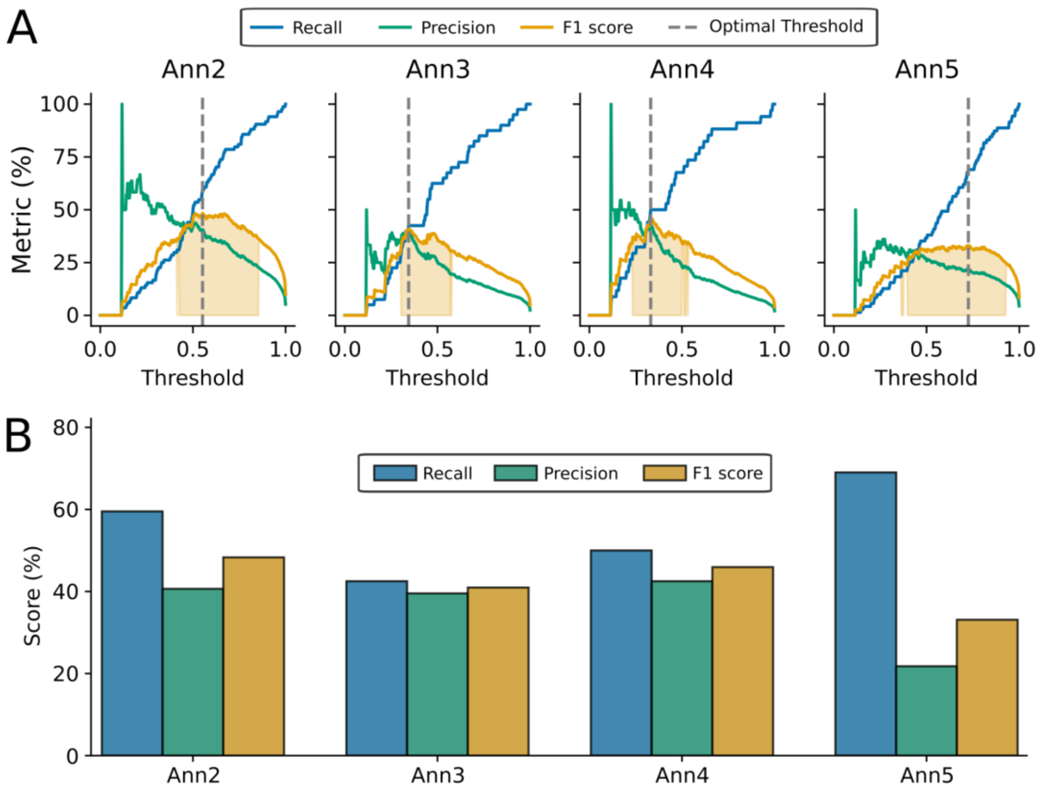


***Supplementary Figure 7. ExoFILT metrics across decision thresholds.***

*(A) Recall, precision, and F1 score for ExoFILT model, for each annotator used as ground truth for performance evaluation (Ann2-Ann5), at all possible decision thresholds. The reported optimal threshold is the one maximizing F1 score for each annotator. Shaded area corresponds to the thresholds where the F1 score stays within 25% of the maximum F1 score.*

*(B) Metrics at the optimal threshold (maximum F1 score) for each ground truth.*

### Supplementary Table 1

| Layer name | Kernel size | Filters / Units |
|---|---|---|
| Conv3D (i) | 1x1x1 | 64 |
| Conv3D (ii) | 1x1x1 | 32 |
| Conv3D (ii') | 3x3x3 | 64 |
| Conv3D (iii) | 1x1x1 | 32 |
| Conv3D (iii') | 5x5x5 | 64 |
| Conv3D (iv) | 1x1x1 | 32 |
| Dense (i) | - | 128 |
| Dense (ii) | - | 64 |
| Dense (iii) | - | 32 |

***Supplementary Table 1. Neural network architecture parameters***

### Supplementary Table 2

| Major class | Ambiguity cause | Configuration of particles | Code |
|---|---|---|---|
| Bona fide | – | Single particle | 1a |
| Bona fide | – | + extra particle | 1b |
| Bona fide | – | + extra cluster | 1c |
| Ambiguous | Superposed objects | + extra particle | 2b |
| Ambiguous | Superposed objects | + extra cluster | 2c |
| Ambiguous | No central intensity | Single particle | 3a |
| Ambiguous | No central intensity | + extra particle | 3b |
| Ambiguous | No central intensity | + extra cluster | 3c |
| Ambiguous | Central particle with low SNR | Single particle | 4a |
| Ambiguous | Central particle with low SNR | + extra particle | 4b |
| Ambiguous | Central particle with low SNR | + extra cluster | 4c |
| Ambiguous | Central particle out of focus | Single particle | 5a |
| Ambiguous | Central particle out of focus | + extra particle | 5b |
| Ambiguous | Central particle out of focus | + extra cluster | 5c |

***Supplementary Table 2. Subclasses of simulated exocytic events.***
*Three subclasses of bona fide events and eleven subclasses of ambiguous events were designed, each of them representing common scenarios usually found in our tracking datasets. The code is defined by a number (shared by ambiguity cause) and a letter (shared by configuration of particles). Note that subclass 2a is not defined, since a single particle cannot be ambiguous due to superposition.*

### Supplementary Table 3

| Ground truth | Evaluating annotator | Recall (%) | Precision (%) | F1 score (%) | Cohen's Kappa |
|---|---|---|---|---|---|
| Ann2 | Ann3 | 36.9 | 77.5 | 50.0 | 0.48 |
| | Ann4 | 34.5 | 85.3 | 49.2 | 0.48 |
| | Ann5 | 34.5 | 40.8 | 37.4 | 0.34 |
| | Average ± SD | 35.3 ± 1.1 | 67.9 ± 19.4 | 45.5 ± 5.7 | 0.43 ± 0.06 |
| Ann3 | Ann2 | 77.5 | 36.9 | 50.0 | 0.48 |
| | Ann4 | 47.5 | 55.9 | 51.4 | 0.50 |
| | Ann5 | 52.5 | 29.6 | 37.8 | 0.36 |
| | Average ± SD | 59.2 ± 13.1 | 40.8 ± 11.1 | 46.4 ± 6.1 | 0.45 ± 0.06 |
| Ann4 | Ann2 | 85.3 | 34.5 | 49.2 | 0.48 |
| | Ann3 | 55.9 | 47.5 | 51.4 | 0.50 |
| | Ann5 | 52.9 | 25.4 | 34.3 | 0.32 |
| | Average ± SD | 64.7 ± 14.6 | 35.8 ± 9.1 | 44.9 ± 7.6 | 0.43 ± 0.08 |
| Ann5 | Ann2 | 40.8 | 34.5 | 37.4 | 0.34 |
| | Ann3 | 29.6 | 52.5 | 37.8 | 0.36 |
| | Ann4 | 25.4 | 52.9 | 34.3 | 0.32 |
| | Average ± SD | 31.9 ± 6.5 | 46.7 ± 8.6 | 36.5 ± 1.6 | 0.34 ± 0.01 |

***Supplementary Table 3. Inter-annotator agreement assessment***

*For each pairwise human-human comparison, recall, precision, F1 score, and Cohen's Kappa were computed by treating one annotator as the ground truth and the other as the prediction, with bona fide events as the positive class. For each ground truth, we also report the mean ± SD of each metric across the evaluated annotators. Note that, since recall and precision depend on the choice of reference, these metrics are exchanged when swapping the roles of the two annotators being compared, while F1 score and Cohen's Kappa remain invariant.*

### Supplementary Table 4

| | Without ExoFILT | With ExoFILT |
|---|---|---|
| Ann2 | 84 | 50 |
| Ann3 | 40 | 17 |
| Ann4 | 34 | 17 |
| Ann5 | 71 | 49 |

***Supplementary Table 4. Number of bona fide events.***

*Summary of bona fide events from $FD_1$ selected by annotators Ann2-Ann5, before and after applying ExoFILT.*

### Supplementary Note 1: Data acquisition

*Saccharomyces cerevisiae* cells (OGY1200) expressing the exocyst subunit Exo84 tagged to three mCherry in tandem (exocyst-mCh) were grown and imaged as explained in Puig-Tintó *et al.*[8]. For the case study, a different strain (OGY1192) from the same study was used, expressing exocyst-mCh and mNeonGreen-Sec1 (mNG-Sec1). All images were collected with a Nikon Ti2-E Eclipse inverted microscope with the following setup: sCMOS Zyla 4.2 camera (Andor, Oxford Instruments), SR HP Apo TIRF 100x/1.49 objective (Nikon), 488 nm and 561 nm lasers (Coherent, OBIS) with iLas2 controlled by Modular V2.0 software (GATACA) and a dual bandpass filter cube (ZET488/561x, ZT488/561rpc and ZET488/561m-TRF as excitation, dichroic and emission filters, respectively). The microscope was controlled by MicroManager software[54]. A cage incubator mounted on the microscope (Okolab) was used to keep a constant 25 °C environment. Simultaneous acquisition was performed with an OptoSplit II Bypass image splitter (Cairn Research).

### Supplementary Note 2: Data preprocessing and automated tracking

The acquired dual-color time-lapse movies were preprocessed with a custom ImageJ (Fiji) script, following a similar approach to what is described by Puig-Tintó *et al.*[8]. First, images were splitted and identified as channel 1 or channel 2. Channel 1 was always exocyst-mCh, while channel 2 data was only used for the case study (mNG-Sec1). The extracellular background was reduced with the rolling ball algorithm in Fiji[55], using a radius of 60 pixels. Afterwards, bleach correction was performed with Fiji *Bleach Correct* plugin (*Exponential Fit*)[56]. Finally, time-lapse movies were further preprocessed with a 3D Gaussian filter, with a radius of 0.5 pixels in the spatial dimensions and 2 frames in the temporal dimension.

A second ImageJ script was developed to perform automated tracking on the exocyst-mCh channel. We performed spot detection and linking with TrackMate[39,40]. Spots were detected with the *Difference of Gaussians* (DoG) method, with a particle diameter of 3 pixels, while linking was done with *Simple LAP tracker*, with a linking maximum distance of 1.5 pixels, a distance gap of 2 pixels, and a frame gap of 1 frame. We excluded puncta located within 10 pixels of the image borders, as well as detections occurring within the first or last 10 frames of the movie. To avoid retaining truncated trajectories resulting from this filtering, we also removed tracks starting before frame 12 or ending after frame N-11 (where N is the total number of frames). This ensured that only tracks fully contained within the analyzed temporal window (i.e., frame 12 to N-11) were retained. Finally, we also eliminated short detections with a track length shorter than 16 frames (~1.8 s), since they were mostly observed to be spurious detections due to low SNR.

### Supplementary Note 3: Permissive filter definition

To develop a filtering strategy for a pre-selection step, we selected nine quantitative features extracted with TrackMate (*Track_Mean_Quality*, *Track_Mean_Q_In*, *Max_Distance_Traveled*, *Track_Displacement*, *Track_Min_Speed*, *Track_Max_Speed*, *Track_Mean_Speed*, *Track_Median_Speed*, *Track_Std_Speed*) together with three additional features previously described[8] (*Track_Quality_Environment*, *Track_Quality_Before*, *Track_Quality_After*), which were specifically designed to detect spatial and temporal overlap with neighboring tracks.

For each of the 12 parameters, we computed the distribution of values among the bona fide events found in $RD_1$. We defined the thresholds of the permissive filter based on the observed minimum and maximum values, extended by 25% on both sides to provide an additional, more safe margin. This empirically defined, permissive filter does not require manual tuning of thresholds to preserve bona fide events in different datasets, while effectively removing clearly ambiguous tracks.

### Supplementary Note 4: Data simulations

Using DeepTrack 2.1[47] and the AnDi-Datasets Python package[48], we integrated multiple parameters to simulate sequences of 10x10-pixels frames mimicking experimental exocytic events, both bona fide and ambiguous (Supplementary Fig. 3). Below, we report the main aspects of these simulations. The combination of different features from our pipeline allowed us to define 3 subclasses of simulated bona fide exocytic events and 11 subclasses of ambiguous events (Supplementary Table 2).

#### *1. Generation of 2D trajectories*

The first step in simulating an exocytic event is defining the trajectories of the particles involved. A central particle, positioned at the center of the simulated area, was always included. Trajectories for additional particles were also defined, either near or far from the central particle, to create ambiguous or bona fide events, respectively (Supplementary Table 2).
All particles were modeled with the AnDi-Datasets Python package as nearly static objects with their centroid confined to a small 2x2 pixel region and a slight subdiffusive movement (fractional Brownian motion with diffusion coefficient of 0.05 pixels$^2$/frame and anomalous exponent of 0.1). The duration of the simulated trajectories was randomly set between 50 and 150 timepoints.

#### *2. Selection of intensity profile*

Exocytic events require multiple copies of the exocyst complex to organize into a sub-diffraction ring-shaped higher-order structure of up to 38 nm of radius[8]. Experimental fluorescent puncta of exocyst-mCh events exhibit a characteristic intensity profile, which reflects the clustering and disassembly of these clusters, often with a variable plateau phase in between. Based on experimental observations, we determined that the intensity profiles of bona fide exocytic events can be approximated by a generalized normal distribution with shape parameter β between 2.5 and 4 ($\beta = 2$ corresponds to a Gaussian; $\beta > 2$ is commonly termed super-Gaussian):

$$f(x) = \frac{\beta}{2\alpha\Gamma(1/\beta)} e^{-(|x-\mu|/\alpha)^{\beta}}$$

where μ is the location parameter, $\alpha > 0$ is the scale parameter, $\beta > 0$ is the shape parameter and Γ denotes the Gamma function.

To introduce more variability, the height and width of the distribution were adjusted to generate asymmetric profiles.

The intensity of additional objects followed a broader range of behaviors (e.g., constant intensity or starting or ending at different times relative to the central particle, etc.). To provide more temporal context, each intensity profile was padded with initial and final frames of zero intensity, representing time before and after the event.

#### *3. Diffraction-limited puncta*

DeepTrack 2.1 was used to generate sequences of diffraction-limited puncta by integrating trajectories, intensity profiles, and realistic imaging conditions. To replicate the experimental setup (Supplementary Note 1), we simulated an optical device for fluorescence imaging with the following parameters: numerical aperture (1.49), wavelength (600 nm), and effective pixel size (129 nm).

With these parameters, a point spread function (PSF) with a full width at half maximum (FWHM) of 245 nm was obtained, resulting in diffraction-limited puncta of approximately 2x2 pixels. In some specific scenarios (Supplementary Table 2, types 5a-5c), we modified the Z value of the central particle to create out of focus spots with a wider PSF, which were then labeled as ambiguous events.

***4. Background noise and SNR***

To accurately mimic heterogeneous exocytic events, it was crucial to simulate not only diffraction-limited spots but also a realistic local environment. For this, we used a small frame size of 10x10 pixels to preserve sufficient local context around the diffraction-limited spots. Since exocytic events occur at the plasma membrane and our imaging system is focused at the equatorial plane of the cell, we modeled an asymmetric background resembling the media-membrane interface, with a higher intracellular signal. Background fluorescence from both the cell and the surrounding media was quantified in experimental data and incorporated into the simulations. The central particle was always positioned close to the simulated membrane, which was randomly oriented.

As mentioned above, additional particles, either isolated or forming bigger clusters, were introduced at varying distances from the central particle to simulate a crowded environment. If the centroid distance between additional and central particles was below 2.5 pixels, the particles were considered overlapping, mimicking ambiguous events.

The signal-to-background ratio of the central particle in bona fide exocytic events was set between 1.15 and 1.3. Some ambiguous events (Supplementary Table 2, types 4a-4c) included central particles with a signal-to-background ratio between 1.05 and 1.075, where intensity fluctuations resembled random noise. In other ambiguous cases (Supplementary Table 2, types 3a-3c), events were generated without a central particle.

***5. Generation of the final simulated dataset***

The final steps for generating the simulated dataset involved adding Poisson noise, globally normalizing each sample, and applying a 3D Gaussian filter for smoothing ($\sigma$ = 1.5 for the temporal dimension, $\sigma$ = 0.5 for spatial dimensions).

By combining a variety of features from the described pipeline, we defined 3 subclasses of simulated bona fide exocytic events and 11 subclasses of ambiguous events (Supplementary Table 2). We generated 20000 videos for each subclass, which provided sufficient simulated data to train our model (training strategy B).

### Supplementary Note 5: Intensity profiles and colocalization of exocyst-mCh and mNG-Sec1

To derive a quantitative timeline of the exocyst-mCh and mNG-Sec1 (Fig. 4A,B), we selected the 98 pairs of bona fide exocyst-mCh and mNG-Sec1 events, and processed their intensity profiles following these steps: 1) For each channel, intensity values along each track were extracted from the raw time-lapse movies; 2) Each intensity profile was corrected by local background subtraction; 3) Each corrected intensity profile was median-normalized; 4) All track pairs were aligned to the end of the exocyst-mCh track; 5) Tracks from each channel were averaged together to obtain the average intensity profile of each protein (Fig. 4A) as well as their corresponding clustering and disassembly times (Fig. 4B).

To compare exocyst-mCh events with different mNG-Sec1 composition (presence or absence) (Fig. 4C and Discussion), we used all exocyst-mCh bona fide events (n = 194) annotated by Ann1. For this analysis we used corrected intensity profiles without applying median normalization. These profiles were smoothed using a Gaussian filter ($\sigma = 3$), and the maximum intensity value of each profile was determined.

Events were further classified based on the 25th percentile of maximum intensity values: tracks in the lowest quartile were defined as dim (n = 49), and the remaining tracks as bright (n = 145). For each category, we computed the fraction of exocyst-mCh events lacking detectable mNG-Sec1 colocalization. Dim exocyst-mCh events were significantly enriched in mNG-Sec1-negative tracks (40.8%, 20/49) compared to bright events (16.6%, 24/145) (odds ratio = 3.48; two-sided Fisher's exact test, $p = 0.0013$).
